# Single cell profiling of the VMH reveals a sexually dimorphic regulatory node of energy expenditure

**DOI:** 10.1101/549725

**Authors:** J. Edward van Veen, Laura G. Kammel, Patricia C. Bunda, Michael Shum, Michelle S. Reid, Jae W. Park, Zhi Zhang, Megan G. Massa, Douglas Arneson, Haley Hrncir, Marc Liesa, Arthur P. Arnold, Xia Yang, Stephanie M. Correa

**Affiliations:** Department of Integrative Biology and Physiology, University of California, Los Angeles, CA, USA; Laboratory of Neuroendocrinology of the Brain Research Institute, University of California, Los Angeles, CA, USA; Molecular, Cellular, and Integrative Physiology Graduate Program, University of California, Los Angeles, CA, USA; Division of Endocrinology, Department of Medicine, and Department of Molecular and Medical Pharmacology, David Geffen School of Medicine, University of California, Los Angeles, CA, USA; Neuroscience Interdepartmental Doctoral Program, University of California, Los Angeles, CA, USA; Molecular Biology Institute, University of California, Los Angeles, CA, USA

## Abstract

Estrogen signaling in the central nervous system promotes weight loss by increasing thermogenesis and physical activity in the ventromedial hypothalamus (VMH), but the precise neuronal populations regulating these aspects of energy expenditure remain unclear. Here we define the molecular and functional heterogeneity of the VMH using single cell RNA sequencing, *in situ* hybridization, chemogenetic activation, and targeted gene knockdown. We describe six molecularly distinct neuron clusters in the VMH. In females, estrogen receptor alpha (ERα) is restricted to neurons expressing tachykinin-1 (*Tac1*) or reprimo (*Rprm*). Further, *Tac1* and *Rprm* expression is enriched in females, a sex difference that is established by permanent effects of gonadal hormones early in life. Finally, while *Tac1* ablation selectively impairs movement, here we show that silencing *Rprm* selectively dysregulates temperature without affecting physical activity. Together this work provides a novel architectural framework whereby distinct and sexually differentiated neuron populations within the VMH mediate sex-specific aspects of metabolic homeostasis.

## Main

Women transitioning to menopause exhibit decreased energy expenditure and decreased fat oxidation compared to age-matched premenopausal women^1^. Similar to humans, rodents exhibit estrogen-induced changes in energy expenditure; female rats exhibit cyclic patterns of wheel running throughout the estrous cycle^2,3^ and female mice exhibit similar cyclicity in temperature and locomotion^4–6^. These effects are mediated by estrogen receptor alpha (ERα) signaling: eliminating ERα either globally or in the central nervous system leads to obesity due increased feeding, reduced movement, and reduced thermogenesis^7–9^. While estrogen-based hormone therapy can improve metabolic profiles after menopause, it is associated with higher cardiovascular disease risk^10^ and, in the case of estrogen plus progestogen therapy, higher breast cancer risk^11^. To ultimately circumvent the risks associated with systemic estrogen therapy, we aim to pinpoint neurons that control systemic energetic balance and define their responses to estrogen signaling.

Recent work has begun to define the neuron populations that mediate the effects of ERα signaling on energy balance. Conditional knockout mouse models suggest that ERα signaling modulates feeding in female mice via neurons of the pro-opiomelanocortin (*Pomc*) lineage, possibly located in the arcuate nucleus (ARC)^5,9,12,13^ or outside the medial basal hypothalamus^14^. Additionally, ERα signaling modulates two types of energy expenditure, spontaneous physical activity and thermogenesis, via neurons of the steroidogenic factor 1 (*Sf1/Nr5a1*) lineage in the ventromedial hypothalamus (VMH)^9,15–17^. However, ERα-expressing neurons of the VMH have many functions. In addition to female-specific roles in energy expenditure, ERα^+^ VMH neurons control fear, territorial aggression, and self defense in males, maternal aggression in females, and mating behaviors in both sexes^18–23^. We hypothesize that these diverse and sex-specific functions are mediated by distinct subpopulations of ERα^+^ neurons. Consistent with this notion, distinct neuronal ensembles are activated in the ventrolateral region of the VMH (VMHvl) of male mice during interactions with male or female conspecifics^18,24^. While these neuron populations remain to be defined, a subset of ERα^+^ neurons in the VMH, which co-express tachykinin 1 (*Tac1*) and oxytocin receptor (*Oxtr*), drive estrogen-dependent changes in physical activity in females^17,25^. However, the VMHvl populations that control other sex-specific behaviors, such as estrogen-dependent increases in thermogenesis, have not been identified.

The VMH is sexually dimorphic with respect to hormone responsiveness, gene expression, neurochemistry, synaptic organization, and neuron function^26–28^. Here, we use RNA sequencing with single cell resolution to test the hypothesis that neurons in this region are heterogenous and sexually dimorphic. We define six major neuron populations in the VMH and a new sexually dimorphic transcript in the VMHvl, reprimo (*Rprm)*, which regulates core body temperature. Collectively, these studies demonstrate that estrogen regulates energy expenditure in females through two intermingled but distinct neuronal subsets, and suggest that the VMH serves as a hormone-responsive nexus of distinct neural circuits controlling metabolic homeostasis.

## RESULTS

### Single Cell Transcriptomics Reveals Neuronal Heterogeneity in the VMH

We used a fluorescent reporter strategy to isolate neurons of the VMH and single cell RNA sequencing to cluster neurons by transcriptional signature. To selectively label VMH neurons, the *Sf1Cre* driver^29^ was crossed to mice carrying a latent allele of tdTomato *(Ai14)*^30^ (Figure 1a). Importantly, this strategy yields tdTomato expression in neurons of the entire VMH upon Cre expression, including in the ventrolateral VMH where it overlaps with ERα immunoreactivity in both males and females (Figure 1b, c) as in ^31^. tdTomato expression in surrounding hypothalamic regions, the dorsomedial hypothalamus (DMH) and the arcuate nucleus (ARC), was detectable but scattered and infrequent (Figure 1b, c, white arrowheads). Fluorescence aided cell sorting (FACS) was performed on single cell suspensions of hypothalami to isolate live neurons of the *Sf1* lineage for single cell transcriptomic analysis (Figure 1d).

**Figure 1.**
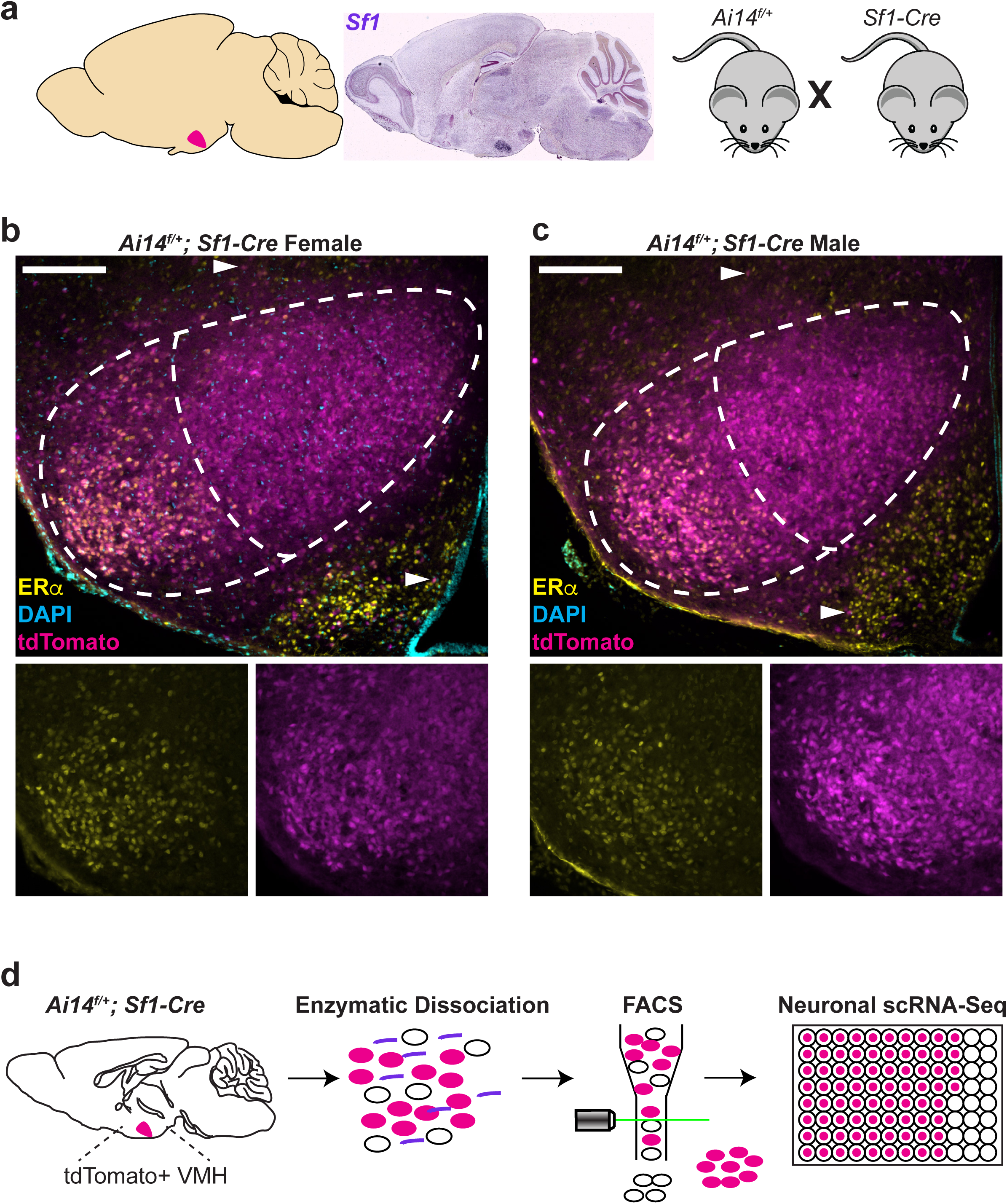
*Sf1* lineage tracing allows for targeted scRNA-seq of the VMH. **a,** In-situ hybridization of *Sf1* transcripts in sagittal section of mouse brain (from Allen Brain Atlas) shows a pattern of expression restricted to the VMH. **b,c,** Mice harboring both the *Sf1-Cre* allele and a latent allele of tdTomato (*Ai14*) show VMH specific fluorescence within the hypothalamus: coronal sections taken from P10 mice, scale bars = 200um. Both female (b) and male (c) VMH show expression of ERα in the VMHvl. As expected, females show higher immunoreactivity. White arrowheads highlight scattered *Sf1* lineage cells outside of the VMH. **d,** Strategy for dissociation followed by FACS and VMH targeted scRNA-seq.

Unicellular transcriptional analysis of 530 single cells from 3 male and 3 female postnatal day (P) 10 mice detected an average (median) of 2556 genes per single cell and revealed strong and consistent expression of the neuronal markers β3-tubulin (*Tubb3*) and neurofilament light peptide (*Nefl*), while very few cells exhibited detectable expression of the glial markers *Gfap* and *Olig1* (Figure 2a). Consistent with the VMH being predominantly glutamatergic, high expression of the glutamatergic marker *Slc17a2* and consistently low expression of the GABAergic marker *Gad2* was observed in all samples (Figure 2a). Finally, to assess how dissociation and FACS sorting may have affected gene expression, we examined immediate early gene expression. Expression of *Fos* and *Arc*, used as a readout for isolation stress and activation^32,33^, appears undetectable in the majority of cells from suspensions obtained from different animals and sexes (Figure 2a). Overall, we conclude that the *Sf1Cre*-mediated fluorescent reporter strategy primarily yields healthy VMH glutamatergic neurons.

**Figure 2.**
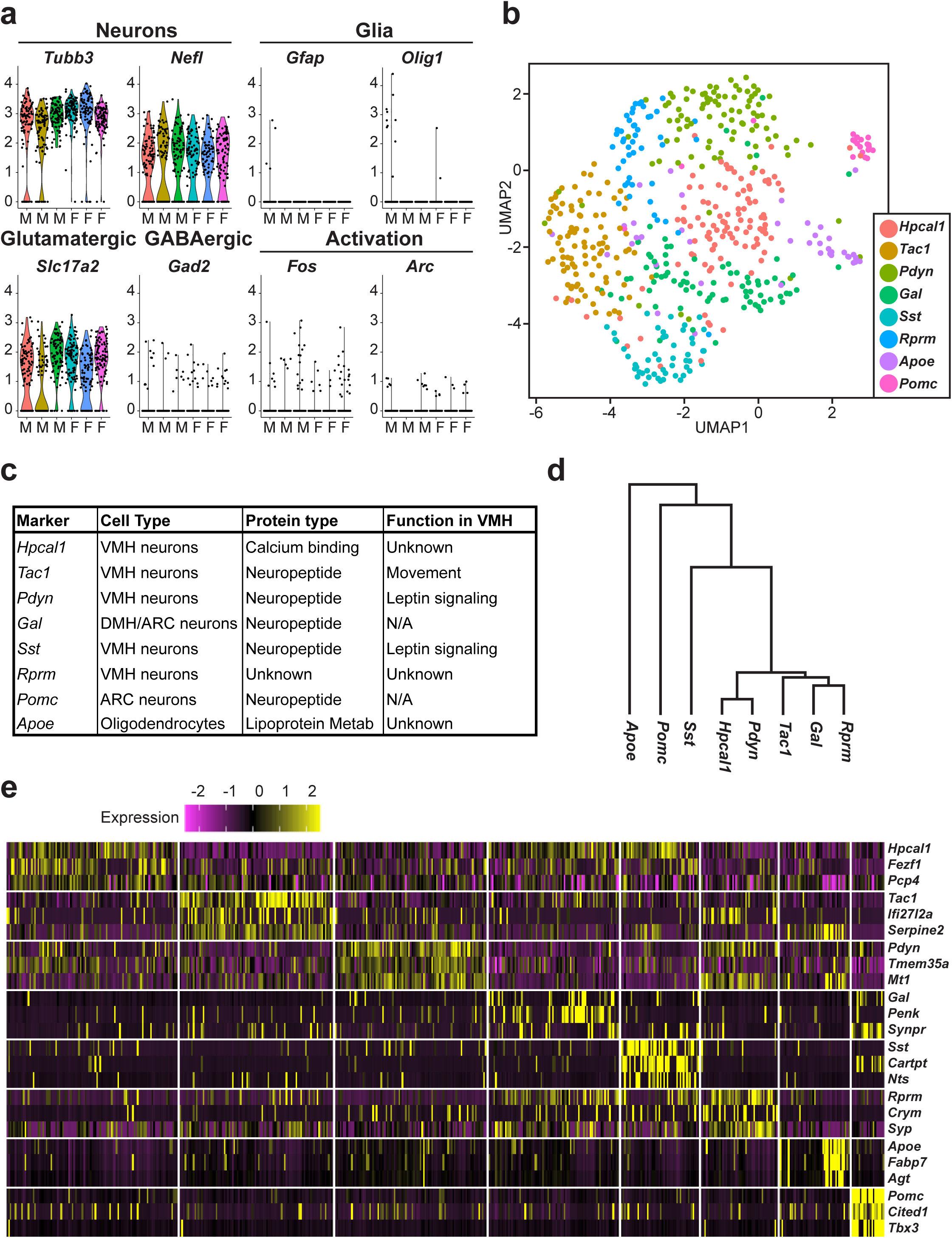
Single cell RNA sequencing reveals non-overlapping gene expression signatures in the VMH. **a,** scRNA sequencing results from (n = 3 female, 3 male) P10 mice showing high expression levels of the neuron specific markers *Tubb3* and *Nefl* with only scattered cells expressing the glial markers *Gfap* and *Olig1*. Cells also express high levels of the glutamatergic marker *Slc17a2*, low levels of the GABAergic marker *Gad2*, and limited expression of the immediate early genes *Fos* and *Arc*. **b,** UMAP showing clusters as defined by marker with highest expression relative to other clusters. **c,** Table showing predicted localization, protein type, and known function of cluster-defining markers. **d,** Hierarchical clustering tree showing relatedness of clusters based on transcriptional signatures. **e,** Heatmap showing expression of top three differentially expressed markers for each cluster.

To determine if VMH neurons show heterogeneity in gene expression profiles, we used a Shared Nearest Neighbor (SNN) algorithm to identify clusters comprised of transcriptionally similar cells^34^. A uniform manifold approximation and projection (UMAP) revealed a main cluster and two divergent clusters (Figure 2b). Within the main cluster, UMAP-based separation was less pronounced, as may be expected when examining neurons of a single transcription factor lineage (*Sf1Cre*). Nevertheless, we detected distinct clusters marked by differential expression of genes that have known and unknown significance in the VMH. We identified a total of eight clusters, hereby identified by the top most differentially expressed transcript within each cluster (Figure 2b, c): *Tac1*, which has been previously demonstrated to promote physical activity in female mice^17^; reprimo (*Rprm*), a TP53 and ERα regulated gene^35^ with no described role in the brain; prodynorphin (*Pdyn*), a gene encoding an endogenous opioid precursor with described roles in leptin-regulated energy homeostasis throughout the hypothalamus^36^; somatostatin (*Sst*), a neuropeptide precursor gene which has hypothalamic roles in the negative regulation of growth hormone axis^37^ and feeding^38^; hippocalcin-like protein 1 (*Hpcal1*) encoding a neuron-specific calcium binding protein; and galanin (*Gal*), a neuropeptide precursor gene shown to increase food consumption when injected into the VMH of rats^39^. In addition to the subpopulations in the principal six clusters, we identified two divergent clusters (Figure 2b, c): a cluster marked by differential expression of proopiomelanocortin (*Pomc*) and many other markers indicating an ARC derived origin, and one marked by differential expression of apolipoprotein E (*Apoe*) and many other markers of glial-like signature.

Comparing overall transcriptional signatures amongst the eight clusters (Figure 2d), the most divergent population are the cells with glial-like signature (*Apoe*^+^), followed by neurons expected to be from the ARC (*Pomc*). Neuron clusters expected to arise from the VMH are more closely related in overall expression signature. Remarkably, the expression of cluster-defining markers appears largely mutually exclusive (Figure 2e), suggesting distinct molecular signatures among neuron clusters of the VMH.

All the neuron clusters identified in the unicellular analysis of the VMH were obtained by analyzing males and females together (Supplementary Figure 1a). We then compared the gene expression profiles between males and females to determine whether sex-specific signatures existed in VMH neurons. The paternally-expressed gene necdin (*Ndn*) had the highest enrichment in males, whereas the proto-oncogene *Araf* had the highest enrichment in females (Supplementary Figure 1b). *Ndn* expression was consistently higher across all clusters in males (Supplementary Figure 1c) and RNA *in situ* hybridization (ISH) confirmed enrichment of *Ndn* transcripts in the male VMH as compared to females (Supplementary Figure 1d). *Araf* expression was detected to be consistently higher in neurons from females across clusters, compared to those from males (Supplementary Figure 1e). ISH of *Araf* was unable to clearly confirm the female biased expression in the VMH, though it appears that *Araf* expression in females might be slightly higher in the VMHvl (Supplementary Figure 1c, d). As *Araf* is a direct effector of the RAF/MEK/ERK MAPK cascade, we sought to determine if this pathway is activated differentially in females. Intriguingly, we found female-specific phosphorylation of MEK in the VMHvl that co-localizes with ERα (Supplementary Figure 1g). This result supports a female specific role of MAPK signaling in the VMHvl as well as *Araf* as a target to be explored to induce changes in energy balance.

### Sex differences established by gonadal hormones

To test the prediction that each neuron cluster generated by gene expression would have a correspondingly distinct spatial distribution within the intact VMH, we detected and localized the expression of the cluster-defining markers using ISH. Further confirming the efficiency of VMH neuron isolation used in the scRNA-seq, expression of all marker genes except *Pomc* was detected within the anatomical boundaries of the VMH (Figure 3 and Supplementary Figure 2a-d). The most restricted expression patterns were observed with *Tac1, Rprm, Pdyn*, and *Sst* (Supplementary Figure 2a-e). Analysis along the rostral-caudal axis revealed sexually dimorphic expression of *Tac1, Rprm*, and *Pdyn* in the caudal VMHvl (Figure 3a-c). Specifically, *Tac1* and *Rprm* expression were both significantly enriched in females within the caudal VMHvl (Figure 3a, b). In contrast, *Pdyn* expression was significantly enriched in males within the caudal VMHvl, although both males and females showed robust expression of *Pdyn* in the dorsomedial VMH (Figure 3c). Finally, we did not detect any major differences in expression of *Sst* between males and females (Figure 3d).

**Figure 3.**
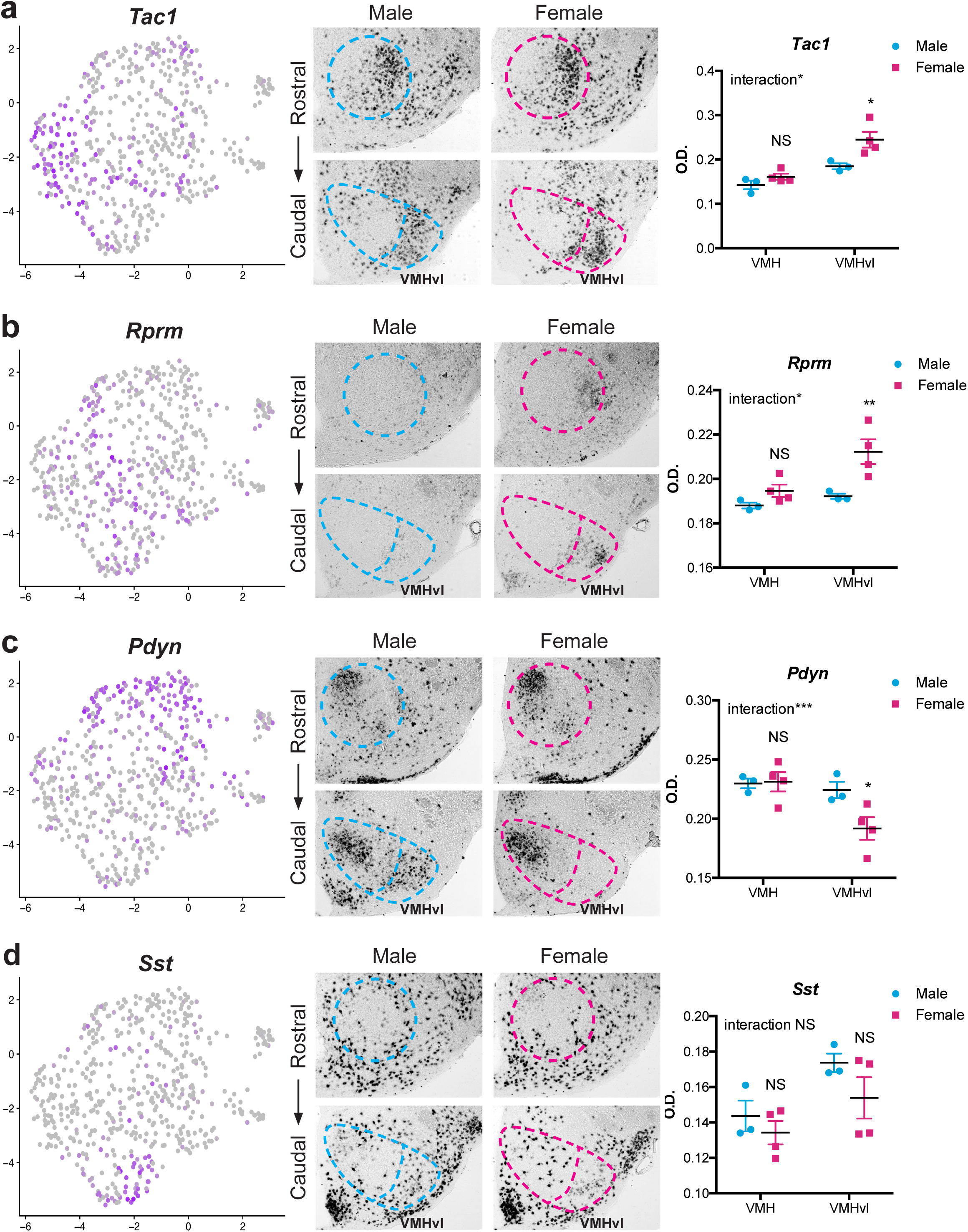
*Tac1, Rprm*, and *Pdyn* are sexually dimorphic genes in the adult VMHvl. Spatial organization of cluster marker within the VMH. Cells positive for **a,** *Tac1*, **b,** *Rprm*, **c,** *Pdyn*, and **d,** *Sst* are identified in purple within the UMAP (left panels). Spatial localization of each cluster marker along the rostral-caudal axis of the VMH in intact males (n = 3 mice) and females (n = 3-4 mice) by chromogenic ISH (right panels). mRNA levels were quantified within the caudal VMH and caudal VMHvl subregion. A statistically significant interaction between sex and ROI was determined for *Tac1* (*F(1,5) = 8.932, p = 0.0305*), *Rprm (F(1,5) = 13.23, p = 0.0149)*, and *Pdyn* (*F(1,5) = 65.84, p = 0.0005*). Post-hoc Sidak’s multiple comparison tests revealed statistically significant sex differences in expression in the caudal VMHvl (*p = 0.0125 for Tac1, p = 0.0071 for Rprm, p = 0.0362 for Pdyn*). Dashed line shows boundary of VMH and VMHvl, in blue for male and magenta for female. Scalebars = 200μm, * = p<0.05, ** = p<0.01, *** = p<0.001.

Sex hormones mediate permanent (organizational) differentiating effects on the brain during development, as well as reversible (activational) effects during adulthood, with additional contributions to sex differences caused by sex chromosome genes expressed within brain cells. To delineate how sexually dimorphic expression of cluster markers develops in the VMHvl, we used the four-core genotypes model^40^ to reveal i) organizational effects of hormones, by comparing XX female vs. XX male or XY male vs. XY female, all gonadectomized (GDX) upon sexual maturity, ii) activational effects of hormones, by comparing GDX vs. intact XX females or XY males, and iii) the effects of sex chromosomes, by comparing XX female vs. XY female or XX male vs. XY male, all GDX, as illustrated in Figure 4a. Expression patterns of both *Tac1* and *Rprm* were unchanged by GDX in females, showing that hormonal activation is not essential for dimorphic expression. Moreover, the presence of the Y chromosome in females did not change *Tac1* or *Rprm* expression, suggesting that the Y chromosome did not have a repressive role on these genes. However, gonadal sex was critical for determining these sexually dimorphic expression patterns, suggesting that these patterns are established during development and are maintained in adulthood (Figure 4b, c). In contrast, the expression pattern of *Pdyn* was distinct between GDX or intact females and gonad-intact males, but not GDX males (Figure 4d, Supplementary Figure 3), suggesting that *Pdyn* expression is maintained by differences in testicular sex hormone signaling in adulthood. Finally, we did not observe any sex differences in *Sst* along any of the three phenotypic comparisons (Figure 4e).

**Figure 4.**
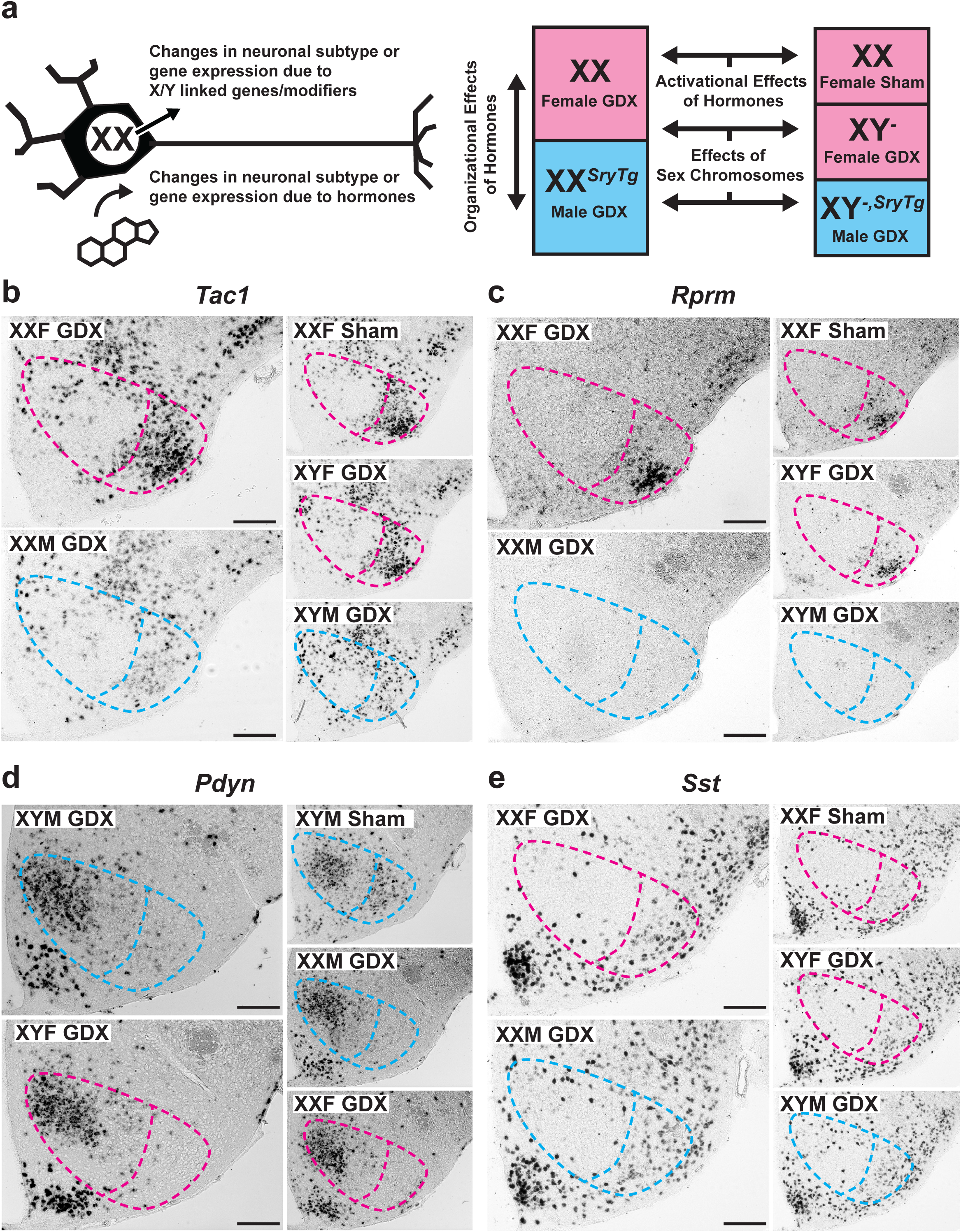
Organizational effects of hormones establish sexual dimorphic expression of cluster markers. **a,** The Four-Core Genotypes (FCG) mouse model, which produces littermates of XX gonadal females (XXF), XX gonadal males (XXM) with an autosomal transgene of the testis-determining gene *Sry* (*Sry Tg*), XY gonadal males (XYM), and XY gonadal females (XYF) with the *Sry Tg*, can be used to determine if the origin of sexually dimorphic gene expression arises due to organizational effects of hormones, activational effects of hormones, or effects due to differences in sex chromosome complement. Expression of **b,** *Tac1*, **c,** *Rprm*, **d,** *Pdyn*, and **e,** *Sst* in the caudal VMH of gonadectomized (GDX) or sham FCG mice (n= 2-3 mice per group) by chromogenic ISH. Dashed line shows boundary of VMH and VMHvl, in blue for male and magenta for female. Scalebars = 200μm.

### Two major estrogen-sensitive populations in the female VMHvl

Estradiol, as a metabolite of testosterone from the testes, plays a major role in the early permanent masculinization of the mammalian brain in males and the expression of male-specific behaviors? To determine if any of the organizational and activational effects of sex hormones on neuronal cluster markers, specifically *Tac1, Rprm*, and *Pdyn*, could be related to estradiol action on ERα? we first examined the expression of ERα by fluorescent ISH (FISH) within each cluster. In females, ERα immunoreactivity was robust in *Tac1*^*+*^ cells and *Rprm*^*+*^ cells (Figure 5a, b), but weak in *Sst*^*+*^ cells (Supplementary Figure 4a). In males, we found that *Pdyn* expression co-localized with ERα immunoreactivity, despite lower ERα expression at both the transcript and protein levels compared to females (Figure 5c, d and Supplementary Figure 3c). We then investigated the spatial relationship of these ERα^+^ neuron populations in females. *Tac1*^*+*^ and *Rprm*^*+*^ cells were highly intermingled within the VMHvl (Figure 5e), but largely spatially distinct from *Sst*^+^ cells (Supplementary Figure 4b, c).

**Figure 5.**
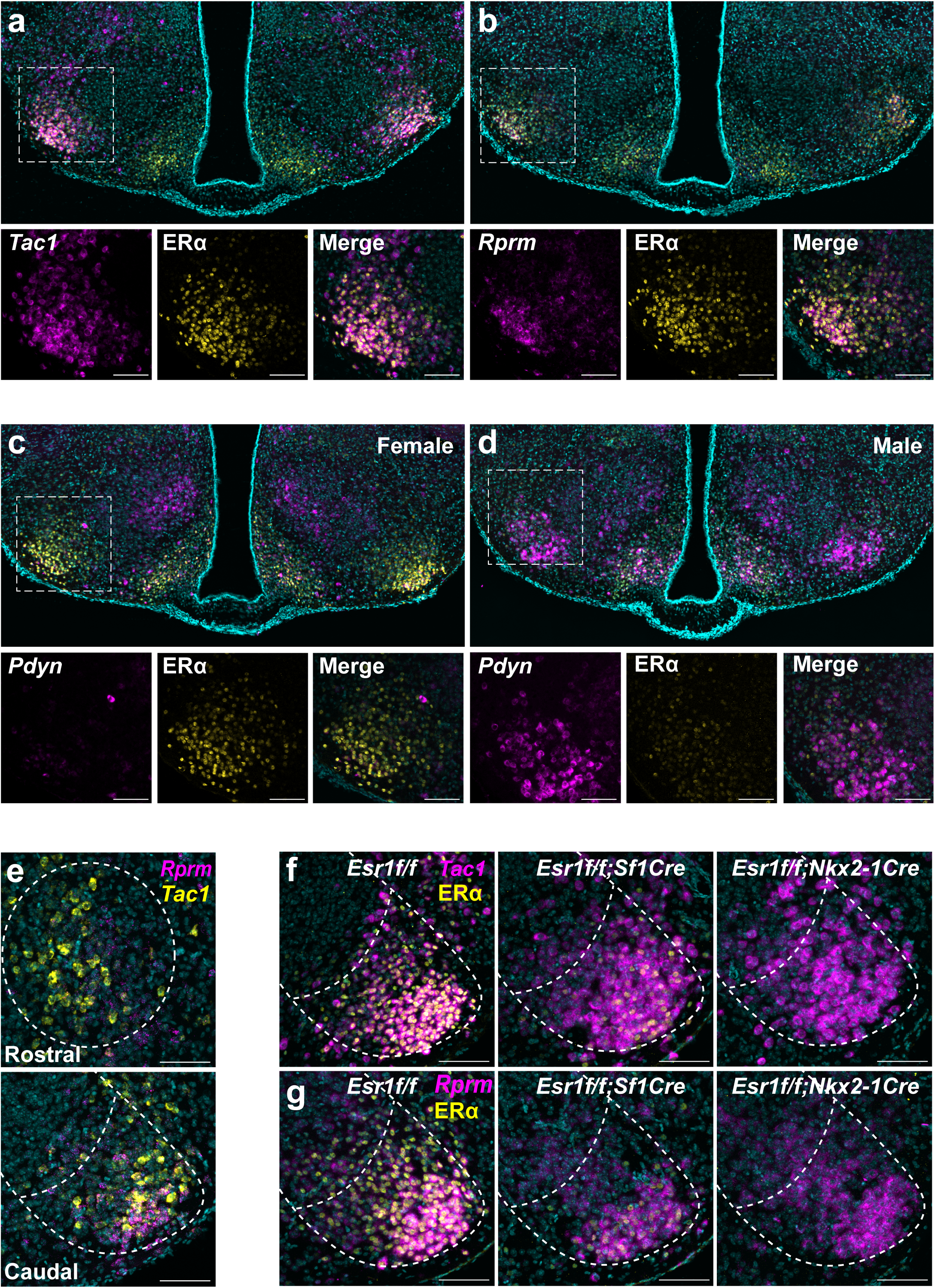
*Tac1*^+^ and *Rprm*^+^ cells are principal ERα-expressing neurons in the female VMHvl. Transcript expression (magenta) of **a,** *Tac1*, and **b,** *Rprm* is shown together with ERa immunoreactivity (yellow) in the VMHvl using fluorescent ISH (FISH, n = 5 female mice). Scalebars on insets = 100μm. Transcript expression of *Pdyn* (magenta) in the VMHvl of **c,** female mice (n = 5) and **d,** male mice (n = 5) and ERα immunoreactivity (yellow) using FISH/IHC. Scalebars on insets = 100μm. **e,** *Rprm* (magenta) and *Tac1* (yellow) transcript expression is visualized using TSA-FISH (n = 5 female mice) in the rostral (top panel) and caudal (bottom panel) VMH. Scalebars = 100μm. Transcript expression (magenta) of **f,** *Tac1* and **g,** *Rprm* together with ERα immunoreactivity (yellow) is visualized in *Esr1*^*f/f*^ mice (n = 6 female mice, left panel), and mice with genetic ablation of ERα in neurons of *Sf1* lineage (*Esr1*^*f/f*^; *Sf1Cre*, n = 4 female mice, middle panel) and in neurons of *Nkx2-1* lineage (*Esr1*^*f/f*^; *Nkx2-1Cre*, n = 3 mice, right panel). Scalebars = 100μm. Images are merged with DAPI (cyan).

Finally, we asked if ERα is required for the sexually dimorphic expression of *Rprm* in females, as previous studies showed that *Tac1* expression was independent of ERα status in the VMHvl. We compared mice with genetic ablation of ERα in the *Sf1* lineage, which results in a substantial loss of ERα^+^ cells in the VMH^9^, and a second mouse model lacking ERα in the *Nkx2-1* lineage, which shows near complete elimination of ERα both in the VMH and ARC^17^. Interestingly, we found that *Rprm* expression, similar to *Tac1* expression, was insensitive to ERα ablation in the female VMHvl (Figure 5f, g). Together, these findings are in line with evidence that early brain masculinization, rather than feminization, is dependent on estradiol-induced regulation of gene expression.

### VMH Expression of Reprimo Regulates the Central Control of Temperature

To discern the role of ERα^+^ VMHvl neurons in energy expenditure, we first chemogenetically activated ERα neurons using the Cre-dependent DREADD (AAV-DIO-hM3Dq^41^) delivered bilaterally to the VMHvl of *Esr1Cre* knockin mice^20^. Administration of the DREADD ligand, clozapine-N-oxide (CNO), elicited a sustained (6 hour) increase in both heat and physical activity in *Esr1Cre* females compared to saline administration in the same mice on a different day and compared to CNO administration to wild-type littermates (Figure 6a, b and Supplementary Figure 5).

**Figure 6.**
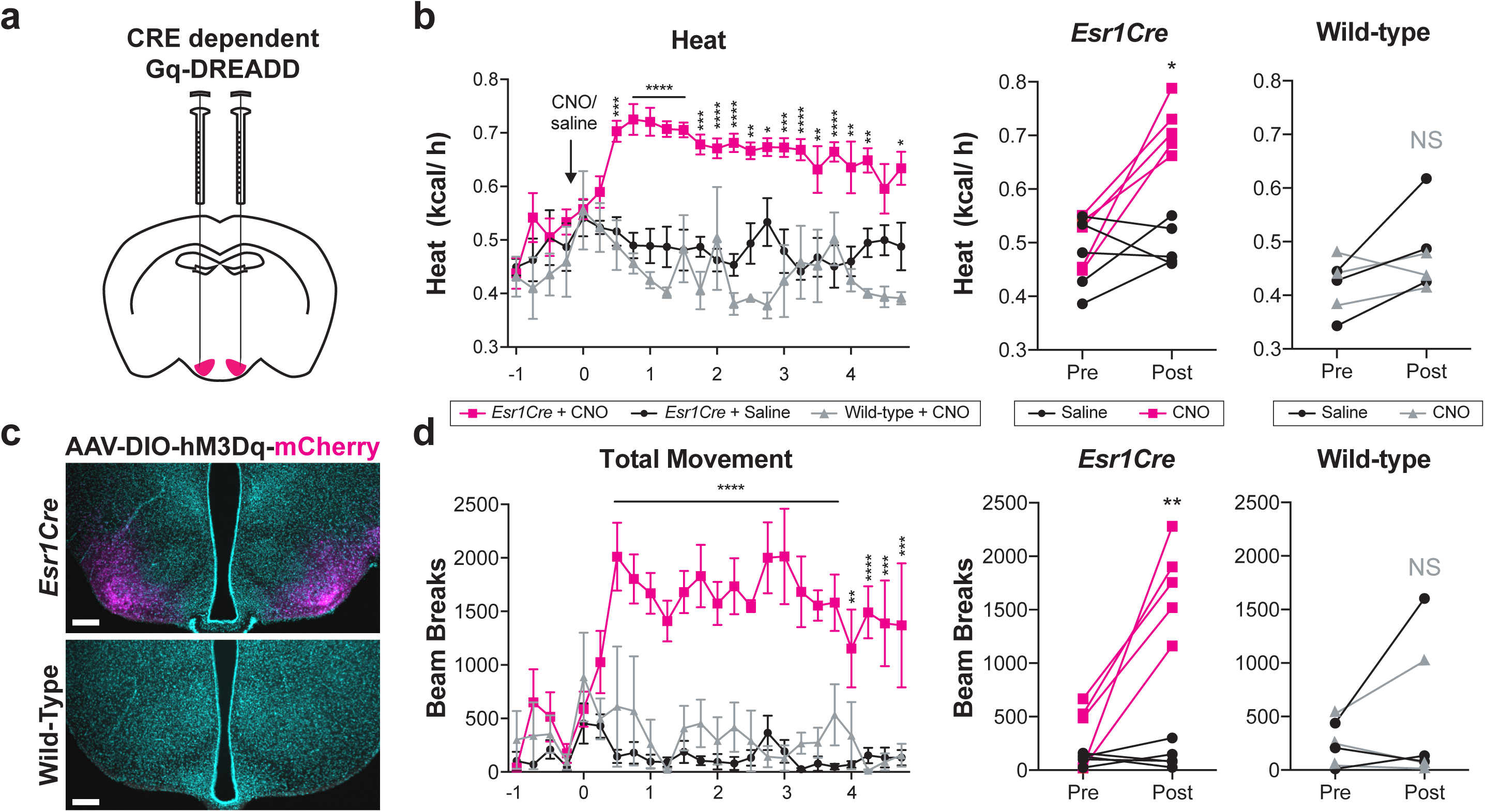
Specific activation of *Esr1*^+^ neurons in the VMHvl causes enhanced movement and thermogenesis. **a,c,** Strategy for and validation of stereotaxic injection of AAVs harboring CRE dependent Gq-coupled DREADDs in the VMHvl **b,** Heat generation increases acutely in *Esr1-Cre* animals after CNO injection but not after saline injection (n = 5 females). Two way RM ANOVA: Time (*F(23,92) = 4.542, p < 0.0001)*, CNO *(F(1,4) = 57.19, p = 0.0016)*, Interaction *(F(23,92) = 3.517, p < 0016).* Wild-type littermate controls stereotaxically injected with the CRE dependent Gq-coupled DREADD show no significant increase in thermogenesis after CNO treatment (n = 3). **d,** Movement increases acutely in *Esr1-Cre* animals after CNO injection but not after saline injection. Two way RM ANOVA: Time (*F(23,92) = 6.361, p < 0.0001)*, CNO *(F(1,4) = 47.17, p = 0.0024)*, Interaction *(F(23,92) = 4.945, p < 0001).* Wild-type animals stereotaxically injected with the CRE dependent Gq-coupled DREADD show no significant increase in movement after CNO treatment. Within-subject changes in average heat and average total movement are shown as averages 0-60 minutes prior to (Pre) and 30-90 minutes following the disturbance to deliver CNO (Post). All subjects were female wild-type mice, ages 10-18 weeks, and singly housed in indirect calorimetry chambers. Posthoc Sidak’s multiple comparison tests were used for pairwise comparisons: * = p<0.05, ** = p<0.01, *** = p<0.001, **** = p<0.0001.

Previous studies suggest that *Tac1*^+^ neurons mediate estrogenic effects on physical activity but not thermogenesis, and that these neurons require *Tac1* expression to fully induce movement^17^. The neuronal population controlling thermogenesis in a sexual dimorphic manner, however, was still elusive. Thus, we hypothesized that the specific role of these newly identified *Rprm*^+^ neurons was to control thermogenesis. To test this hypothesis, we silenced *Rprm* gene function within the VMHvl using cell permeable siRNA pools delivered via bilateral stereotaxic injections (Figure 7a). When compared to animals injected with a non-targeting siRNA, the animals injected with *Rprm* targeting siRNA showed a significant increase in body temperature (Figure 7b, c). The increase in temperature was persistent across time points (Figure 7b) and was significant both in the active night phase and the inactive day phase (Figure 7c).

**Figure 7.**
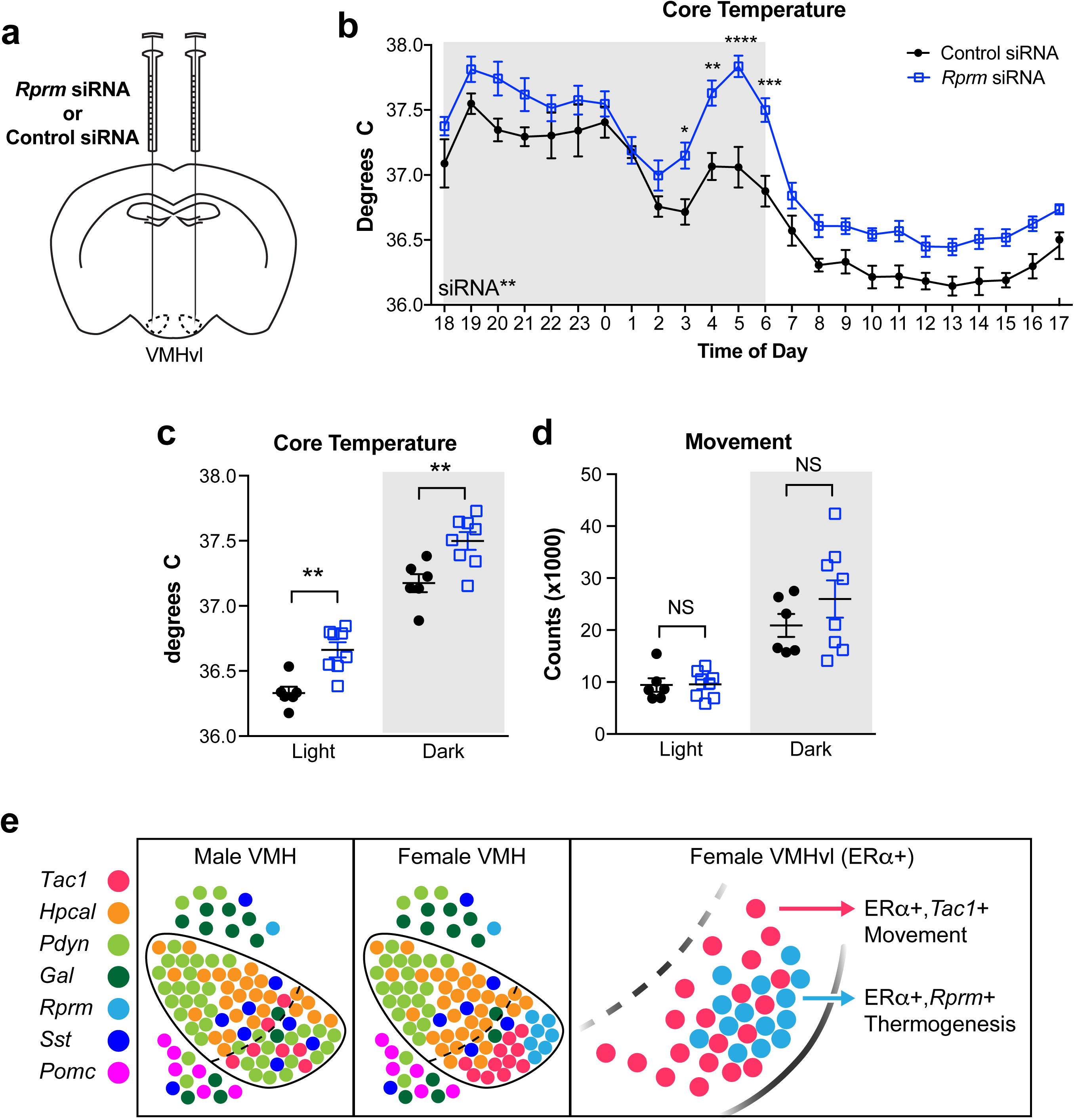
Temperature is dysregulated in mice lacking *Rprm*. **a,** Strategy for stereotaxic injection of cell-permeable siRNA pools either targeting *Rprm* (n = 6) or non-targeting (n = 8). **b,** Core temperature is significantly increased in animals injected with *Rprm* targeting siRNA pools compared to animals injected with non-targeting siRNA pools. Two way RM ANOVA: Interaction (*F(23,276) = 1.653, p = 0.0329)*, Time *(F(23,276) = 67.31, p <0.0001)*, siRNA *(F(1, 12) = 18.31, p = 0011)*. **c,** The effect of *Rprm* depletion on core temperature is significant in both the sleep (day) phase and active (night) phase compared to non-targeting controls. Two way RM ANOVA: Interaction (*F(1,12) = 1.653, p = 0.9408)*, Time *(F(1,12) = 330.1, p <0.0001)*, siRNA *(F(1, 12) = 18.31, p = 0011)*. **d,** *Rprm* depletion showed no significant effect on movement in either sleep or active phase when compared to non-targeting controls. All subjects were female wild-type mice, ages 10-20 weeks, and singly housed. **e,** diagram of neuronal populations found in the VMH with focus on sexually dimorphic female VMHvl. Posthoc Sidak’s multiple comparison tests were used for pairwise comparisons: * = p<0.05, ** = p<0.01, *** = p<0.001, **** = p<0.0001.

Importantly, *Rprm* knockdown did not induce changes in physical activity (Figure 7d), further supporting the notion that *Tac1*^*+*^ neuronal function does not overlap with *Rprm*^*+*^ neuronal function. These data demonstrate for the first time that there are at least two classes of ERα^+^ neurons in the VMHvl that are functionally distinct, and together coordinate the female specific effects on physical activity and thermogenesis of this hormone-responsive region.

## DISCUSSION

This study used the power of single cell RNA sequencing as a starting point for a high resolution atlas of the VMH, one of the brain’s longest known sexually dimorphic regions. By performing our sequencing ten days after birth, we were able to discover populations established in development, before they are altered by hormonal changes associated with experience or puberty. Extensive validation experiments confirm all of the populations identified by sequencing analysis. Additionally, *in situ* hybridization comparing the sexes revealed novel sexually dimorphic populations in the caudal VMHvl that scRNA sequencing did not resolve. We detected only scattered reads from *Esr1* in our sequencing, but combining immunohistochemistry with in situ hybridization clearly identified specific populations that overlap with ERα expression, notably including the sexually dimorphic subpopulations of the VMH. Finally, *in vivo* validation experiments confirm that ERα^+^ neurons in the VMHvl play unique roles in regulating energy expenditure in females, and extend this finding by functionally and molecularly subdividing these neurons. Functional dissection of sex differences in the neural circuits that control food intake and energy expenditure is critical to understanding the biological basis of gender differences in the control of body weight.

Single cell RNA sequencing has led to dramatic improvements in understanding diverse cell populations of the hypothalamus, using dissociated brain tissue without purification of cell types ^42,43^. Here, we extend these studies by using mice harboring a genetically encoded fluorescent lineage tracer (*Sf1-Cre; Ai14*). This approach was tailored to allow specific purification of VMH neurons by FACS prior to scRNA-Seq. We identified six major and two minor clusters of cells with distinct transcriptional signatures. Notably, previous studies do not identify the two clusters marked *Hpcal1* or *Rprm*, nor do they assign *Tac1, Pdyn, Sst*, or *Gal* populations to the VMH ^42,43^. In contrast, targeted profiling of the *Sf1* lineage allows the study of the VMH with unprecedented resolution.

As the overall anatomy of the VMH is conserved between males and females (Figure 1b-c, 2b, Supplementary Figure 1a), it is curious how activation of equivalent neurons can evoke sex specific behaviors. It is now generally accepted that sexually dimorphic behaviors are sexually differentiated by sex hormones during critical developmental periods (reviewed in ^44,45^). Estrogen signaling in the VMH has clearly demonstrated roles in coordinating the increased movement and thermogenesis that accompany the sexually receptive period in female mice^16,46^. Indeed, partial ablation of *Esr1*^*+*^ neurons in the VMH impairs BAT thermogenesis in females^9^. Additionally, activation of neurons in the VMHvl increases physical activity in females, but not in males, in a manner that is dependent on both circulating estrogens and hypothalamic expression of ERα^17^. Here, we find that specific activation of *Esr1*^*+*^ neurons in female VMHvl increases physical activity and body temperature, supporting the notion that *Esr1*^*+*^ VMHvl neurons coordinate estrogen-dependent energy expenditure.

The present results suggest that sex hormone signaling during development drives the emergence of two female-specific subpopulations of *Esr1*^*+*^ neurons, defined by largely mutually exclusive expression of either *Tac1* or *Rprm*. Previous studies link the *Tac1*^*+*^ subset of ERα^+^ neurons to the regulation of physical activity in females, without affecting thermogenesis^17^. To date, several lines of evidence have also implicated estrogen signaling in the VMH, primarily dependent on inhibition of AMP-kinase (AMPK), in enhanced BAT thermogenesis through activation of the sympathetic nervous system (SNS)^16,47^. We now report that silencing *Rprm* function selectively alters temperature without significantly affecting movement. Together, our expression analyses and functional studies suggest a model in which estrogens act on the *Tac1* and *Rprm* neuron clusters to increase energy expenditure in females by two distinct mechanisms (Figure 7e). The intermingling of *Tac1*^+^ and *Rprm*^+^ neurons within the VMHvl is intriguing. In the future, it will be very interesting to examine whether *Rprm*^+^ neurons and *Tac1*^+^ neurons exhibit crosstalk and how their circuit wiring differs, but this will require currently unavailable tools, such as a *Rprm-Cre* mouse, to allow for specific labeling, activation, and inhibition of these populations.

These studies also uncovered a male-specific pattern of expression in the VMHvl, defined by the expression of *Pdyn*. A notable difference between the *Pdyn* and *Tac1* or *Rprm* subpopulations is that maintenance of *Pdyn* in the VMHvl expression requires circulating testicular hormones. This result is made more striking by the observation that *Pdyn* expression in the dorsomedial VMH (VMHdm) is contrastingly unaffected by castration. Dynorphins, the products of *Pdyn*, are potent endogenous opioid peptides^48^ with demonstrated roles in reward, addiction, and stress^49^. It will be very interesting in future studies to examine the role of VMH *Pdyn* expression in male behavior, and how these behaviors might be modified by circulating levels of testosterone. In addition to the observed sub-specialization of the VMHvl, we observed a limited number of VMH-wide sex differences in gene expression, including male-biased expression of *Ndn*, which appears to be due to fewer *Ndn*^+^ cells in the VMH of females compared to males.

Entry into menopause is associated with significant increases in visceral abdominal fat and body weight. Surprisingly, this is not associated with an increase in caloric intake. Instead, adiposity correlates with a decrease in overall energy expenditure, which manifests most strikingly during sleep^1^, implying that an increasingly sedentary lifestyle cannot be the primary determinant. Because postmenopausal obesity confers enhanced risks of cardiovascular disease and breast cancer, there is a clear and urgent need to find new strategies to combat weight gain. Replacing hormones lost during menopause, such as estrogen and progesterone, can bring about weight loss, but these treatments themselves carry potential cardiovascular and cancer risks. We speculate that *Tac1*^+^ and *Rprm*^*+*^ neurons are important nodes in the dysregulation of energy expenditure accompanying the abrupt decline in circulating sex hormones experienced during menopause. As such, the molecular mechanism whereby these neurons control thermogenesis will be of interest for the treatment of post-menopausal obesity.

## Experimental Procedures

### Mice

All studies were carried out in accordance with the recommendations in the Guide for the Care and Use of Laboratory Animals of the National Institutes of Health. UCLA is AALAS accredited and the UCLA Institutional Animal Care and Use Committee (IACUC) approved all animal procedures. Mice expressing the *Sf1Cre* driver transgene (*Tg(Nr5a1-cre)7Lowl*), the *Nkx2-1Cre* driver transgene (*Tg(Nkx2-1-cre)2Sand*), and the *Ai14-*tdTomato reporter with loxP-flanked STOP cassette (*Gt(ROSA)26Sort*^*m14(CAG-tdTomato)Hze*^) were maintained on a C57BL/6 genetic background. The *Esr1 floxed* allele (*Esr1*^*tm1Sakh*^) was maintained on a CD-1;129P2 mixed background. Breeder male “Four Core Genotypes” mice (FCG, background C57BL/6J) possess a Y chromosome deleted for the testis-determining gene *Sry*, and an *Sry* transgene inserted into chromosome 3. The four genotypes include XX and XY gonadal males (XXM and XYM), and XX and XY gonadal females (XXF and XYF). Genotypes were discriminated using genomic PCR as described in ^50^. All other experiments were carried out on C57BL/6J mice (JAX 000664). Except for gonadectomy studies, all experiments were performed in intact males and intact cycling females.

### scRNA sequencing

We labeled all VMH neurons by crossing the Cre-dependent tdTomato reporter (*Ai14*)^30^ to the *Sf1Cre* driver^29^ to generate *Ai14*^*f/f*^; *Sf1Cre* mice. Because the tdTomato signal is largely restricted to the VMH, a fairly large hypothalamic region was collected under fluorescent illumination. Cells were dissociated using a papain-based enzymatic process (Worthington Biochemical). VMH neurons were sorted by FACS using parameters that select for tdTomato signal. Because tdTomato is expressed in processes and projections, we enriched for cell bodies using a nuclear DNA dye (cell permeant DRAQ5, ThermoFisher). Dead cells were excluded by eliminating cells stained by NucBlue (cell impermeant DAPI). Doublet discrimination was used to ensure single cells were deposited into each well. Individual tdTomato^+^ neurons were sorted into each well of a 96-well plate (Precise WTA kits, BD). The Precise WTA single cell sequencing kits include a well index to identify each cell and a unique molecular index (UMI) to identify each transcript and reduce bias due to PCR amplification. Libraries were prepared according to manufacturer’s instructions and sequenced on an Illumina NextSeq 500 using paired end 2 x 75 bp mode.

Expression data were analyzed using the R package Seurat^51^. Normalized data were scaled with a linear regression model based on number of unique molecular identifiers (UMIs) per cell and percentage of reads from the mitochondrial genome to remove unwanted sources of variability and to normalize gene expression data. Analyses included all genes expressed in ≥ 2 cells, and all cells expressing ≥ 500 genes and a fraction of mitochondrial reads < 0.17. To cluster cells based on transcriptome similarity, we used Shared Nearest Neighbor (SNN) algorithm^52^. For each cell cluster, marker genes were determined by comparing expression in the given cluster against all other clusters using the smart local moving algorithm to iteratively group clusters together^52^. To determine sex differences, all female neurons passing initial filtering were compared to all male neurons passing initial filtering.

### Mouse Procedures

Mice were anaesthetized with isofluorane and received analgesics (0.01mg/mL buprenorphine, 0.58mg/mL carprofen) pre- and post-surgery. Bilateral ovariectomy and castration surgery with complete removal of the ovaries or the testes was performed on adult mice. For Figure 4, sham or gonadectomized control mice (XXF and XYM) and gonadectomized FCG mice from separate experimental batches are shown together. The Cre-dependent AAV8-hM3Dq-mCherry DREADD (Addgene, titer ≥ 4×10^12^ vg/mL, 200 nL to each side) was injected bilaterally into the VMHvl of adult female mice (coordinates: A-P: −1.56 mm from Bregma; lateral: ±0.85 mm from Bregma; D-V: 5.6 mm from the cortex). After 2 weeks of recovery, mice received i.p. injections of CNO (0.3 mg/kg) or vehicle (saline, 0.15%DMSO) 3 hr after the onset of the light phase. Saline and CNO were administered on consecutive days in a randomized balanced design. siRNA pools against *Rprm* or non-targeting controls (Dharmacon, 0.4 mM, 350 nL to each side) were delivered to the VMHvl as described above. Indirect calorimetry was performed in Oxymax metabolic chambers (Columbus Instruments). Gross movement and core body temperature were measured using an IP-implanted G2 eMitter and VitalView software (Starr Life Sciences).

### RNA probe generation

Digoxigenin (DIG)- or fluorescein (FITC)-labeled sense and antisense riboprobes for somatostatin (*Sst)*, reprimo (*Rprm)*, tachykinin 1 (*Tac1)*, prodynorphin (*Pdyn)*, necdin (*Ndn)*, and proto-oncogene, serine/threonine kinase A-Raf (*Araf)* were in vitro transcribed from template cDNA using T7/T3/SP6 RNA polymerase (DIG/FITC RNA labeling kit, Roche) and purified with RNA Clean & Concentrator (Zymo Research). For template cDNA generation, PCR products for individual genes were amplified from a hypothalamic cDNA library and cloned into pCR 2.1-TOPO or pCR II-TOPO (Invitrogen) for all probes except *Tac1*, which was previously described^17^. Plasmid DNA was isolated from bacterial cultures using ZymoPURE II Plasmid Midiprep kit (Zymo Research), linearized by restriction digest, and purified with DNA Clean & Concentrator (Zymo Research). All PCR products, except *Araf*, were generated using reference primer sequences from the Allen Brain Institute. For *Araf*, cDNA was generated from bases 639-942 (NM_009703.2).

### In situ hybridization

The ISH protocol was partially adapted from previously published methods^17^. 18μm-thick cryosections containing the VMH were obtained from paraformaldehyde-fixed mouse brains. Day 1: Upon defrosting to room temperature (rt), slides were washed in PBS, postfixed in 4%PFA, and washed again. TSA-fluorescent ISH (FISH) slides were also incubated in 3%H_2_O_2_ for 30min to quench endogenous peroxidase activity. To permeabilize the tissues, slides were incubated in proteinase K (1ug/mL) for TSA-FISH and chromogenic ISH (CISH), or 0.3%Triton X-100 in PBS for combined FISH-IHC. CISH slides were postfixed again in 4%PFA for 5 min. Slides were incubated in hyb solution containing probe overnight at 65C. Day 2: Coverslips were removed in solution containing 5x SSC heated to 65C. Slides were then subject to a series of stringency washes, then blocked in NTT containing 2%blocking reagent and HISS for 2 h at rt. Slides were incubated in antibody solution containing either anti-DIG-AP (1:5,000), anti-FITC-AP (1:5,000), or anti-DIG-POD (1:4,000) in 4C overnight. FISH-IHC slides were additionally incubated in anti-ERα (Rb, 1:1000). Day 3: Slides were washed in NTT, then NTTML (0.15M NaCl, 0.1M Tris pH 9.5, 50mM MgCl_2_, 2mM levamisole, and 0.1%Tween-20) to quench endogenous phosphatase activity. Slides were developed in INT/BCIP solution (Roche). FISH-IHC slides were blocked in 10%normal goat serum for 1hr at rt, and incubated with anti-rabbit 647 for 2 h at rt, and incubated in HNPP/Fast red working solution according to manufacturer’s instructions (HNPP Fluorescent Detection Set, Roche). To stop the reaction, the slides were washed 3x 5min in PBS, counterstained with DAPI, and immediately stored in −20C to prevent HNPP/Fastred diffusion. TSA-FISH slides were incubated in working solution containing Cy5 Plus tyramide according to manufacturer’s instructions (Perkin Elmer). Slides were then washed in NTT and incubated in 3%H_2_O_2_ for 30min to quench the first tyramide reaction. Slides were then washed 3x 5min in NTT, blocked in in NTT containing 2%blocking reagent and HISS for 2 h at rt, and incubated overnight in anti-FITC-POD (1:4,000). Day 4: TSA-FISH slides were washed in NTT, and incubated in working solution containing FITC Plus tyramide according to manufacturer’s instructions (Perkin Elmer). The reaction was terminated with NTT and slides were counterstained in DAPI. Control experiments using sense riboprobes and no probes showed negligible signal. Additionally, performing the TSA reaction following 3%H_2_O_2_ for 30min in the absence of a second POD incubation confirmed adequate quenching. Probes with weaker signal intensity were developed first in TSA-FISH.

### Image Acquisition and Quantification

All CISH experiments with imaged in brightfield on a DM1000 LED microscope (Leica) using 5X or 10X objectives. Semi-quantitative optical density (O.D.) measurements of mRNA in CISH slides were obtained with ImageJ (NIH) following calibration with a calibrated step tablet (Kodak), according to standard protocols^53^. Measurements from the left and right VMH were averaged to calculate the mean O.D. for each animal using predetermined ROIs based on the Franklin and Paxinos Mouse Brain Atlas. Sex differences in O.D. between the caudal VMH and caudal VMHvl were determined by two-way ANOVA with Bonferroni multiple-comparison correction. FISH and IHC experiments were imaged on a LSM780 confocal microscope (Zeiss) using 10X or 20X objectives. Tile-scanned images were stitched using Zen Black (Zeiss). All images were taken with the same z-sampling interval for a given objective and z-stacks were merged to obtain maximum intensity projections. Cyan/magenta/yellow pseudo-colors were applied to all fluorescent images for accessibility. Image processing, limited to brightness and contrast, was performed using the Leica Application Suite (Leica), Zen Black (Zeiss), ImageJ (NIH), and Photoshop (Adobe).

## Acknowledgements

The research was supported by UCLA Division of Life Sciences funds to SMC, NIH K01 DK098320 to SMC, NIH UL1TR001811 and Iris Cantor-UCLA Women’s Health Center/UCLA National Center of Excellence in Women’s Health Pilot Awards to SMC and ZZ, UCSD/UCLA Diabetes Research Center NIH P30 DK063491 Pilot and Feasibility awards to SMC and ML, NIH grants DK104363 and NS103088 to XY, NIH grants HD076125 and HL131182 to APA, UCLA Department of Medicine Chair commitment to ML, pre-doctoral NRSA (F31 AG051381) and Hyde Fellowship to LGK, UCLA Dissertation Year Fellowships to LGK and DA, Canadian Diabetes Association Postdoctoral fellowship to MS, American Heart Association Postdoctoral Fellowship (18POST33960457) to ZZ, and NSF Graduate Research Fellowship to MGM. The authors thank Carolina De La Cruz for technical assistance.

## Author Contributions

JEV, LGK, and SMC conceived of and designed the studies. JEV, LGK, PCB, MS, MSR, JWP, ZZ, MGM, HH, and SMC acquired and analyzed data. JEV, LGK, PCB, MS, DA, ML, APA, XY, and SMC contributed to data interpretation. JEV, LGK, and SMC wrote the manuscript with substantial input from MS, ZZ, MGM, DA, ML, APA, and XY.

