## Supplemental Figures for "Single cell profiling of the VMH reveals a sexually dimorphic regulatory node of energy expenditure"

Supplementary Figures:

**Supplementary Figure 1.** The overall architecture of the VMH is conserved between males and females.

**Supplementary Figure 2.** Clustering and expression of non-specific markers and markers outside of the VMH.

**Supplementary Figure 3.** Sexually dimorphic expression of *Pdyn* in the VMHvl is not maintained by differences in ovarian sex hormone signaling in adulthood.

**Supplementary Figure 4.** *Sst*[+] cells show limited ER$\alpha$ immunoreactivity in the female VMHvl.

**Supplementary Figure 5.** DREADD activation increases cFOS immunoreactivity.

### van Veen, Kammel, et al. Fig. S1

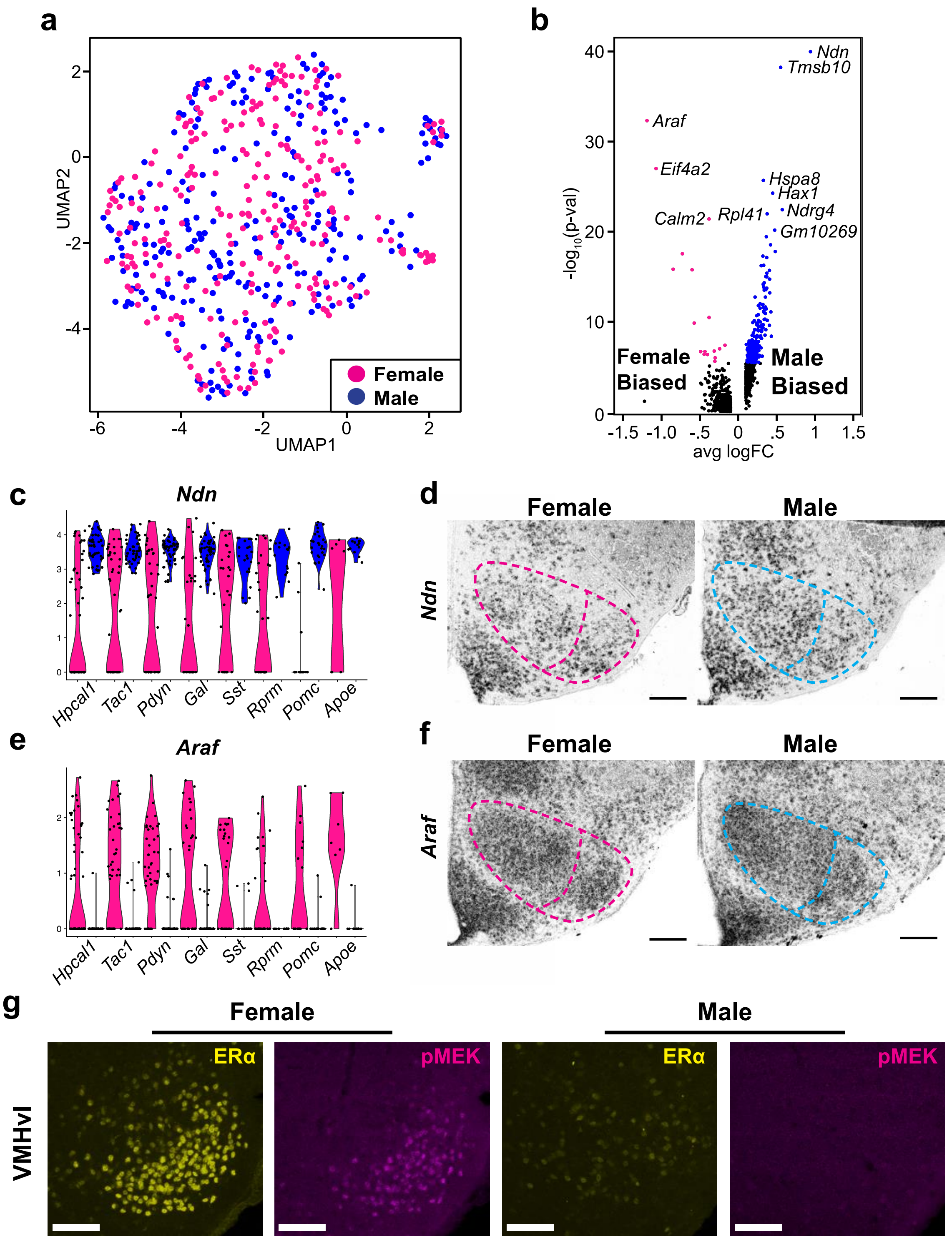


22

**Supplementary Figure 1. The overall architecture of the VMH is conserved between males and females. Related to Figure 2. a,** UMAP showing that male and female neurons are present in all clusters identified. **b,** Volcano plot showing differences in gene expression when comparing male and female VMH transcriptomes. Female biased (Adj. $P < .05$) genes shown in pink, male biased (Adj. $P < .05$) shown in blue. **c,** Violin plots showing male biased expression of *Ndn* across all VMH clusters. **d,** ISH validating male biased expression of *Ndn* in the VMH, scalebars = 200µm. **e,** Violin plots showing female biased expression of *Araf* across all clusters identified. **f,** ISH showing expression of *Araf* in male and female VMH, scalebars = 200µm. **g,** immunofluorescence demonstrating strong pMEK immunoreactivity in a subset of ER$\alpha$ expressing female neurons in the VMHvl, but no observable pMEK staining in the male VMHvl, scalebars = 100µm.

van Veen, Kammel, et al. Fig. S2

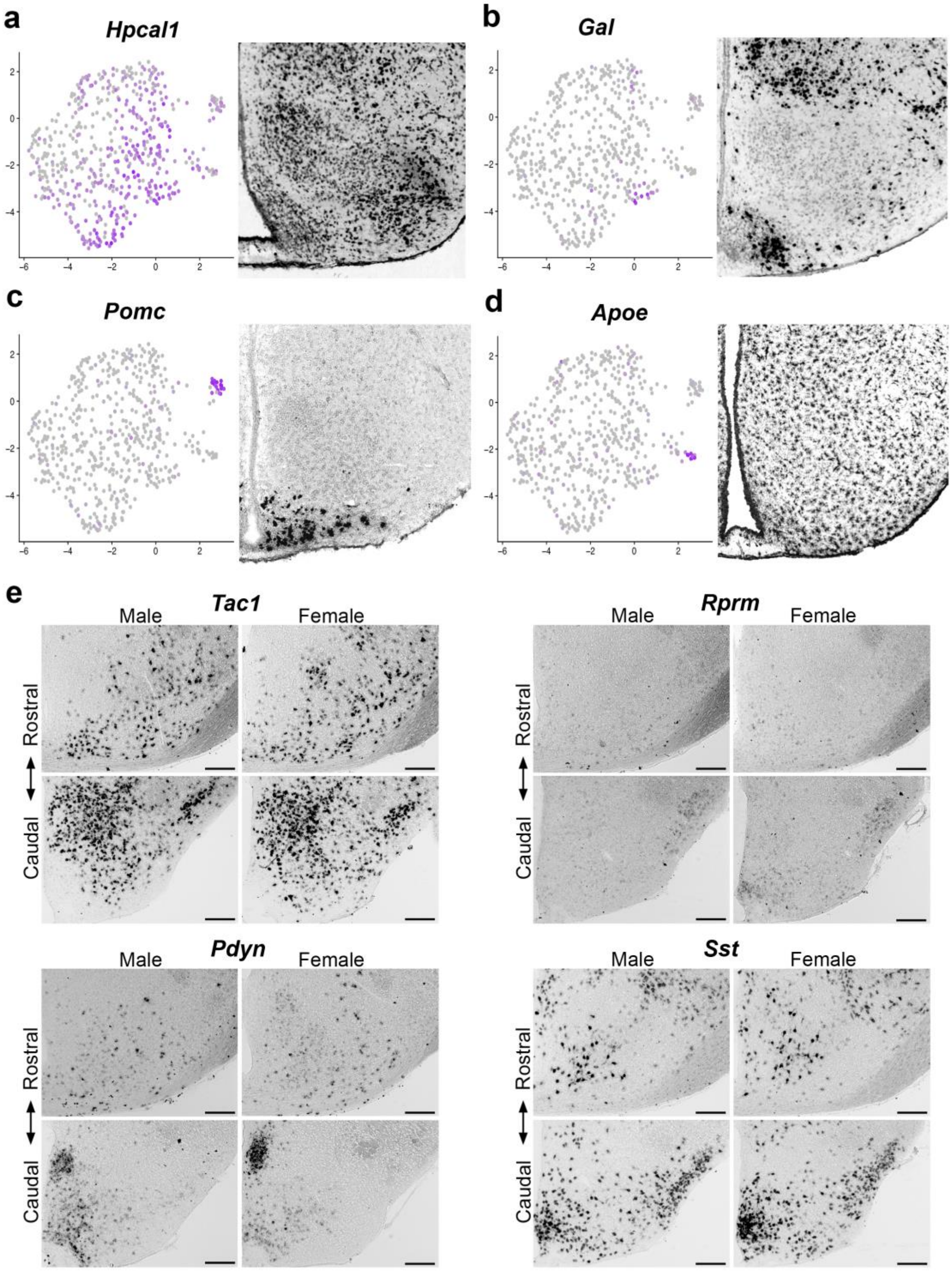


36

**Supplementary Figure 2. Clustering and expression of non-specific markers and markers outside of the VMH. Related to Figure 3. a,** *Hpcal1* expression appears diffusely in the UMAP clustering analysis, and similarly, ISH shows diffuse *Hpcal1* expression in the VMH. **b,** *Gal* expression is restricted to only a handful of cells on the UMAP, and similarly, ISH shows *Gal* expression in scattered VMH cells. **c,** *Pomc* expression is strongest outside of the central UMAP, and similarly, ISH shows *Pomc* expression strongest in the ARC. **d,** *Apoe* expression is strongest outside of the central UMAP. ISH shows *Apoe* expression with no particular pattern. **e,** expression of *Tac1*, *Rprm*, *Pdyn*, and *Sst* in brain areas adjacent to the VMH along the rostral-caudal axis in males (n=3) and females (n=4). Scalebars = 200μm. Images in **a-d** from Allen Brain Atlas.

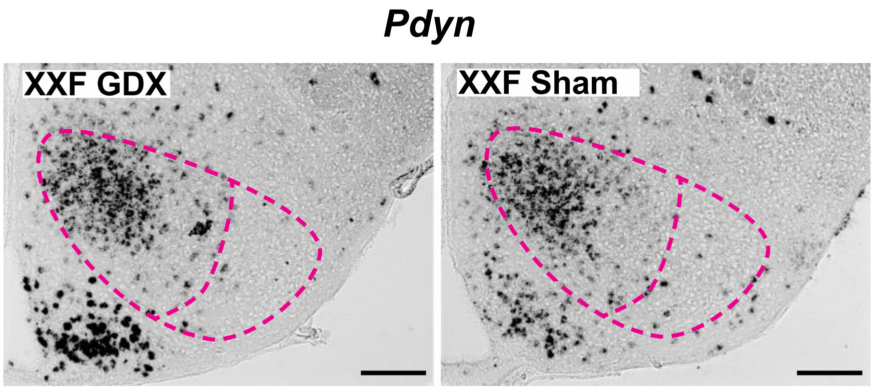


**Supplementary Figure 3. Sexually dimorphic expression of *Pdyn* in the VMHvl is not maintained by differences in ovarian sex hormone signaling in adulthood. Related to Figure 4.** Expression of *Pdyn* in the caudal VMH of gonadectomized (GDX) or sham XXF FCG mice (n=3 mice) by chromogenic ISH. Dashed line shows boundary of VMH and VMHvl, in magenta. Scalebars = 200μm.

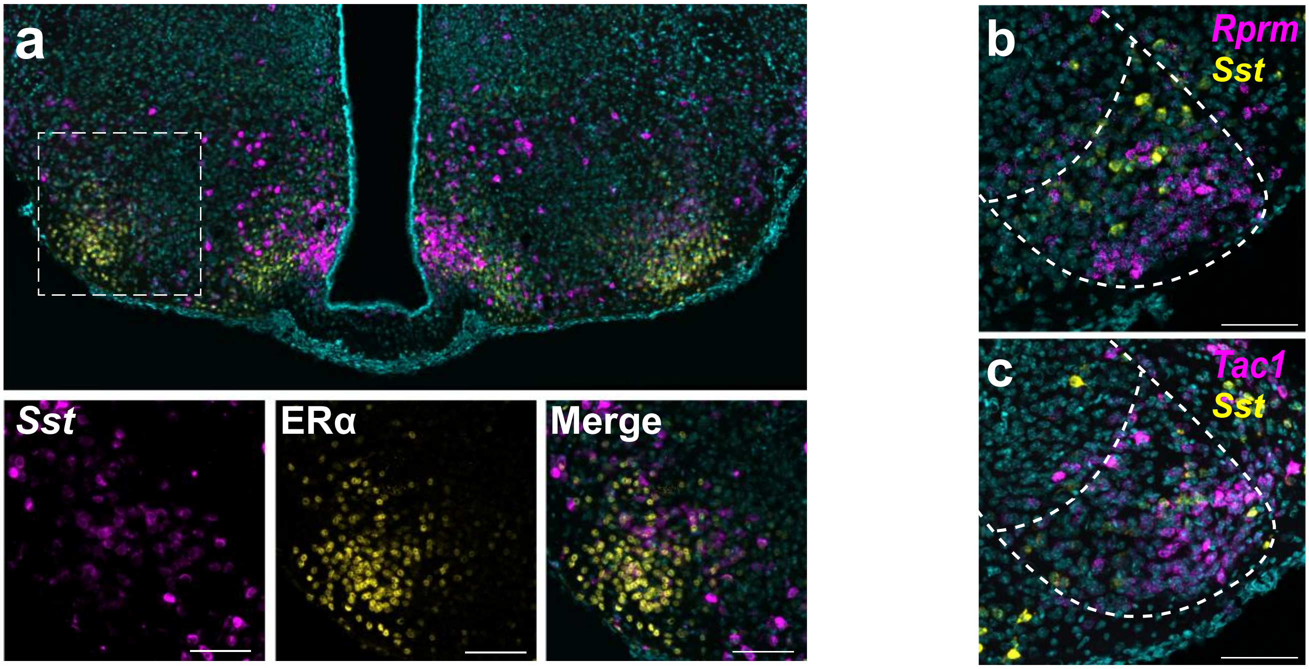


**Supplementary Figure 4. *Sst*$^{+}$ cells show limited ERα immunoreactivity in the female VMHvl. Related to Figure 5. a,** Transcript expression of *Sst* (magenta) is shown together with ERα immunoreactivity (yellow) in the VMHvl using fluorescent ISH (FISH, n=5 female mice). Scalebars on insets = 100µm. Transcript expression (magenta) of **b,** *Rprm*, and **c,** *Tac1* is visualized with *Sst* transcript expression (yellow) using TSA-FISH (n=5 female mice) in the caudal VMH. Scalebars = 100µm. Images are merged with DAPI (cyan).

**van Veen, Kammel, et al. Fig. S5**

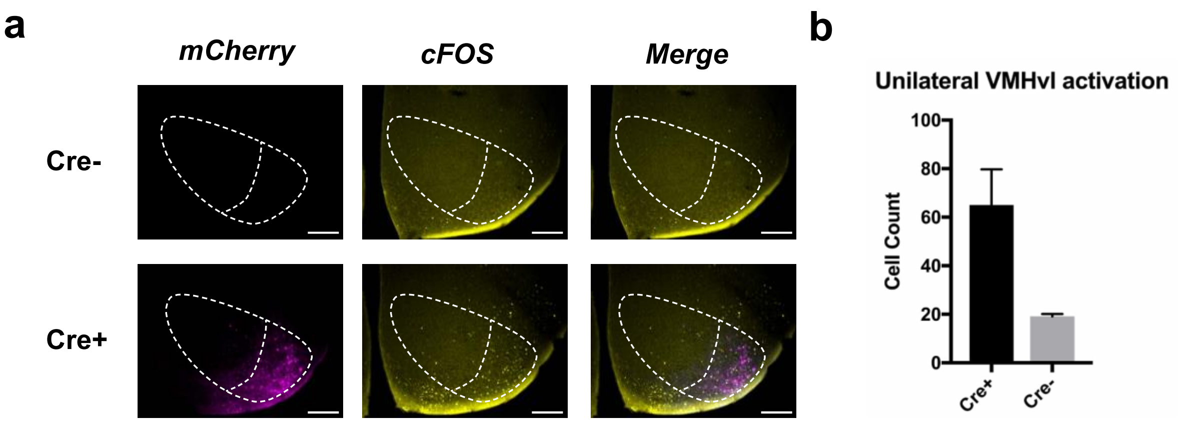


**Supplementary Figure 5. DREADD activation increases cFOS immunoreactivity. Related to Figure 6. a,** Representative images of increased mCherry (magenta) and cFOS (yellow) in the VMHvl of *Esr1Cre* (Cre+, n = 5) mice compared to wild-type (Cre-, n = 3) injected with a viral, cre-dependent mCherry-fused Gq-coupled DREADD. **b,** cFOS immunoreactivity in the VMHvl (quantified on one side per animal) is increased in Cre+ subjects 90 minutes following CNO injection. Scalebars = 200μm.
